# Stochastic Simulation Algorithm for effective spreading dynamics on Time-evolving Adaptive NetworX (SSATAN-X)

**DOI:** 10.1101/2021.11.22.469498

**Authors:** Nadezdha Malysheva, Max von Kleist

## Abstract

Modelling and simulating the dynamics of pathogen spreading has been proven crucial to inform public heath decisions, containment strategies, as well as cost-effectiveness calculations. Pathogen spreading is often modelled as a stochastic process that is driven by pathogen exposure on time-evolving contact networks. In *adaptive networks*, the spreading process depends not only on the dynamics of a contact network, but vice versa, infection dynamics may alter risk behaviour and thus feed back onto contact dynamics, leading to emergent complex dynamics. However, stochastic simulation of pathogen spreading processes on *adaptive networks* is currently computationally prohibitive.

In this manuscript, we propose SSATAN-X, a new algorithm for the accurate stochastic simulation of pathogen spreading on *adaptive networks*. The key idea of SSATAN-X is to only capture the contact dynamics that are relevant to the spreading process. We show that SSATAN-X captures the contact dynamics and consequently the spreading dynamics accurately. The algorithm achieves up to 100 fold speed-up over the state-of-art stochastic simulation algorithm (SSA). The speed-up with SSATAN-X further increases when the contact dynamics are fast in relation to the spreading process, i.e. if contacts are short-lived and per-exposure infection risks are small, as applicable to most infectious diseases.

We envision that SSATAN-X may extend the scope of analysis of pathogen spreading on *adaptive networks*. Moreover, it may serve to create benchmark data sets to validate novel numerical approaches for simulation, or for the data-driven analysis of the spreading dynamics on *adaptive networks*. A C++ implementation of the algorithm is available at https://github.com/nmalysheva/SSATAN-X.

**Author summary:** Modelling and simulating of infectious disease spreading supports public heath decisions, such as prevention and containment strategies and allows to perform cost-effectiveness calculations. Detailed modelling approaches consider stochastic pathogen spreading on time-evolving contact networks. In *adaptive networks*, the spreading process depends not only on the dynamics of a contact network, but vice versa, infection dynamics may alter risk behaviour and thus feed back onto contact dynamics.

Stochastic simulation of these complex dynamics is currently computationally prohibitive.

We propose a new algorithm (SSATAN-X) that can significantly speed up stochastic simulations on *adaptive networks*, while being accurate at the same time. Our algorithm achieves this speed-up by only considering the contact dynamics that are relevant to the spreading process. The benefit of algorithm is particularly pronounced when contacts are short-lived and per-exposure infection risks are small, which is applicable to most infectious diseases.

We envision that SSATAN-X may allow simulation and analysis of pathogen spreading on more complex *adaptive networks* than previously possible. Moreover, data sets may be created with SSATAN-X that are useful for benchmarking novel numerical schemes and analytic approaches.

## Introduction

Modelling and simulation of contagion spreading to forecast disease prevalence and to assess different public health interventions has a long history in mathematical biology [1]: Traditional approaches involve compartmental models, such as susceptible-infectious-recovered (SIR) or susceptible-infected-susceptible (SIS) [2]. However, traditional compartmental models do not capture relevant spreading paths and may even provide incorrect predictions [3]. Network models incorporate more realism, explicitly considering interactions (=edges) between individuals and locations (=nodes). Within this context, mostly static networks have been studied in the past [4, 5]. However, in static networks, the extent of linkage of nodes is determined only after integrating information derived from observing the contact (or spreading) process over a period of time. Several examples highlight that analysis of static networks lacks important temporal information about causal paths that underlie the spreading process, consequently yielding false conclusions for the control of the spreading process [6]. In order to capture the temporal causality of the underlying system, different time-evolving network models have been introduced recently [7, 8]. A particularly relevant class are adaptive networks. In this class of network models, the network structure itself changes dynamically in response to the dynamics of the spreading process [9, 10]. Examples are the concurrency of sexual partnerships which is thought to be important for HIV spread [11–13] or measures of self-isolation (quarantine) for individuals diagnosed with SARS-CoV2. This creates a feedback loop between the spreading dynamics *on the network* and the dynamics *of the network* itself, leading to emergent complex behaviour. Epidemic spreading has been extensively studied on these types of networks to understand the influence of social contact structure on disease prevalence [14–17]. However, the dynamics of contagion spreading on adaptive networks are usually complex, and analytical results can only be obtained in special cases [17, 18]. This makes numerical studies based on stochastic simulations indispensable.

Several different approaches are available to date: The stochastic simulation algorithm (SSA) [19] allows for the exact numerical simulation of contagion spreading on adaptive networks, as used in [20–22]. Ready-to-use implementations are available in the network modelling packages Largenet2 [23] and EpiFire [24]. A major drawback of the SSA is that it may generate computational overhead, when the contact dynamics are considerably faster than the contagion dynamics, i.e. if only a tiny fraction of contacts result in contagion transmission. While the per-contact transmission probability can be of the order 10^−1^ some airborne diseases like influenza [25], it is usually much lower in sexually transmitted diseases like HIV (order 10^−3^ – 10^−2^) [26, 27]. The computational overhead becomes even more pronounced, when the outcome of intervention strategies, like face masks, vaccination, prophylaxis or ‘treatment as prevention’ are studied which further lower the per-contact contagion transmission probability [28–31].

Two types of algorithms are used in this context: (i) Inexact methods discretize the time and then perform parallel updates of the contact structure and the contagion spreading (akin to a tau-leaping procedure in systems biology [32, 33]). Time discretization and synchronous updates are implemented in EpiModel [34] and NepidemiX (http://nepidemix.irmacs.sfu.ca/). (ii) The method proposed by Vestergaard [35] borrows ideas from the ‘integral method’ originally proposed and applied in the Systems Biology field [36]. It assumes deterministic contact dynamics, e.g. in Vestergaard [35], a temporal set of static contact configurations is used. If the contact dynamics would in fact be deterministic for the given time intervals, the method then estimates the exact time to the next spreading event. However, in Vestergaard’s original implementation the underlying contact network is not *adaptive* any longer; i.e., the network configurations affect the spreading dynamics, but not *vice versa*. Therefore, this approach is limited to particular scenarios, in which human behaviour is not adapted as the pandemic unfolds, in contrast to what is currently observed for the SARS-CoV2 outbreak.

In this article be develop an efficient and exact stochastic simulation method that allows to predict contagion spreading in the case when both the contact dynamics, as well as the spreading dynamics are governed by inter-dependent stochastic processes, i.e. human behaviour is adapted as a function of the epidemic process. The algorithm is written in C++, available through https://github.com/nmalysheva/SSATAN-X

## Materials and methods

### Model: spreading process on an adaptive contact network

In the following, we introduce the proposed simulation algorithm using an example of a simple spreading process that evolves on a time-dependent, adaptive contact network.

Consider a population Ω = {**S**, **I**, **D**} that consists of susceptible- **S**, infected- **I** and diagnosed individuals **D**. Each of these individuals has a number of contacts that change over time. Infection (e.g. transmission of a virus) is possible from an infected to a susceptible, or from a diagnosed to a susceptible individual, only if the respective individuals are in contact. For an airborne pathogen, e.g. influenza or SARS-CoV2, a ‘contact’ denotes a period of time where individuals are in close proximity (e.g. within the same room), whereas for sexually transmitted infections it denotes a sexual relationship.

The contacts change over time. To describe the contact dynamics of the model, each individual is assigned rates of gaining and loosing contacts over time.

Moreover, in case the individual is diagnosed with the infection, he/she may become aware of his/her status, which affects the individual’s behaviour. In case of our modelling exercise, the individual cuts all of his/her contacts, and the individual rate of establishing new contact drops to 30% of its pre-diagnosis value. Thus, this simple model describes adaptive dynamics, where the contact dynamics change depending on the epidemic state, Fig. 1C.

**Fig 1.**
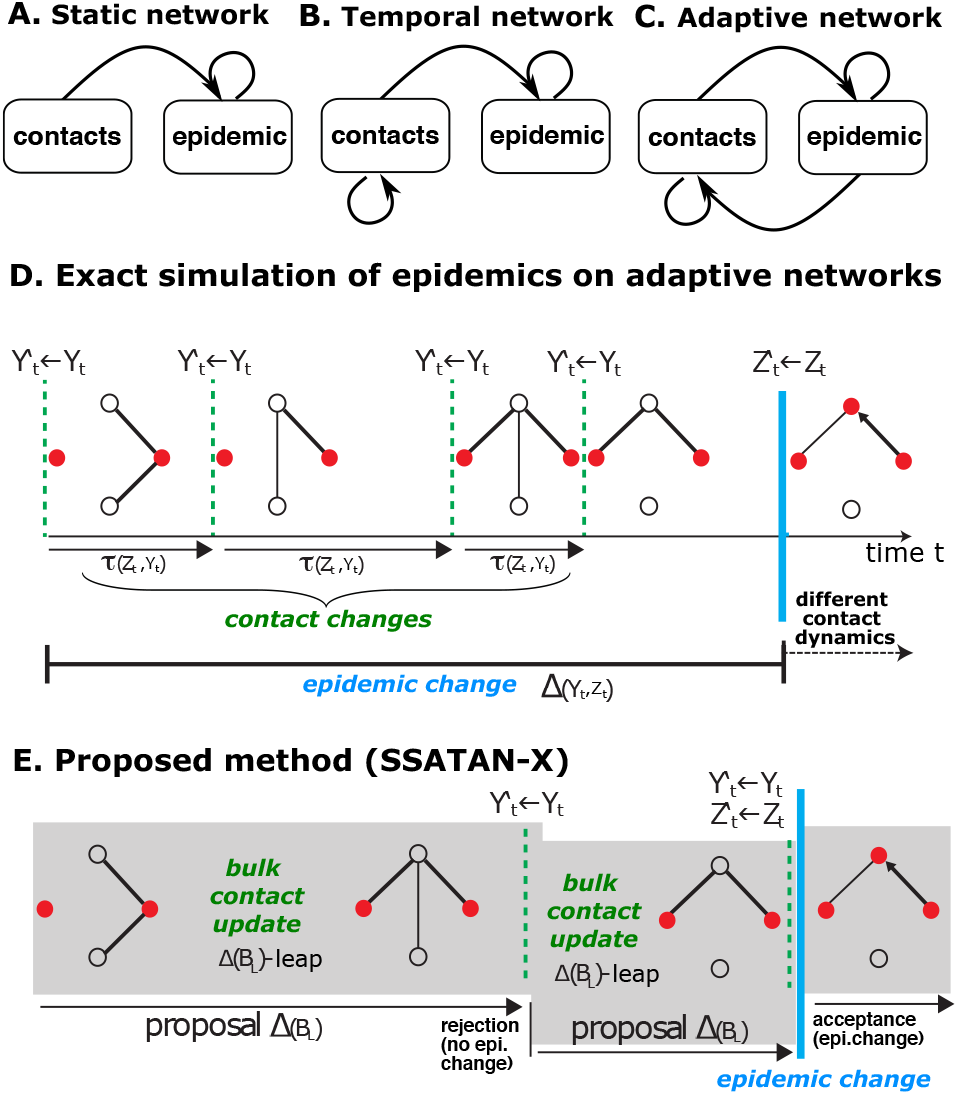
Methods for simulating spreading processes on networks. We consider two processes *Y*|*Z*: *E* → *E* and *Z*|*Y*: *V* → *V*. The first process *Y*|*Z* affects the edges *E* of the network (topology, *contact dynamics*), while the second process *Z*|*Y* affects the state of the nodes *V* (*epidemic dynamics*). **A-C**. Different approaches for modelling spreading processes on networks. **A.** In static network approaches, the network topology *E*, which determines the linkage and thus the spreading process *Z*, remains fixed. **B.** In temporal networks, the network topology evolves according to some *contact dynamics Y*: *E* → *E*, which is independent of the epidemic process *Z*|*Y*. **C.** Adaptive networks denote a class of co-evolving stochastic processes, where both the contact structure affects the spreading process Z |Y, and the spreading process alters the contact dynamics *Y*|*Z*. **D.** Exact stochastic simulation [19] of spreading processes on adaptive networks. In the stochastic simulation algorithm, every change to the network is explicitly modelled, including changes in the edges/topology, as well as changes to the state of the nodes (empty circles vs. filled red circles). **E.** Proposed method: The proposed method only considers the *net* effect of topological changes on the *epidemic dynamics*. An upper bound *B_L_* is computed such that *B_L_* ≥ ∑_*j*∈*Z*_*a_j_*(*V, E*), and the network topology is bulk updated for the time step Δ(*B_L_*) ~ exp(*B_L_*) using *τ*-leaping. A change to the state of the nodes (*epidemic process*) is conducted if *r* ≤ *a*_0,*Z*_/*B_L_*; 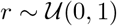 and *a*_0_, *z* = ∑_*j*∈*Z*_*a_j_*(*V, E*).

The presented model can be described as an undirected weighted temporal *Contact Network G*(*t*) = {*E*(*t*), *V*(*t*)}, where

- *V* denotes the set of nodes {*v*_1_(*t*), *v*_2_(*t*), …, *v_N_*(*t*)} ∈ Ω (the infected, diagnosed and susceptible individuals). Each node *v_i_* ∈ *V* has the following individual characteristics: (i) a rate at which a new contact is made *λ_i_* and (ii) a rate of loosing an existing contact *θ_i_*.
- The set of edges is denoted *E*(*t*) = {*e*_(*j,k*)_(*t*)} with *e_j,k_* = (*v_j_, v_k_; γ*), *j* ≠ *k*, if *v_j_* and *v_k_* are in contact. Parameter *γ* represents the transmission rate from *j* to *k*, which is > 0 if node *j* is infected or diagnosed and node *k* is susceptible and otherwise 0.

We consider a continuous-time Markov process that evolves the contact network *G*(*t*) in time. The possible events that can occur on the network can be classified into two main groups:

1. Contact dynamics 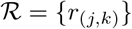:

- Assembling of a new contact. In our example, for each pair of nodes (*v_j_, v_k_*), *j* ≠ *k* which is not connected by an edge, the rate of assembling an edge is defined by *λ*_(*j,k*)_ = *λ_j_* · *λ_k_* i.e. product of the assembling rates of the two nodes.
- Disassembling of an existing contact. For each pair of nodes (*v_j_, v_k_*), *j* ≠ *k* that are connected by an edge, the rate of disassembling is defined as *θ*_(*j,k*)_ = *θ_j_* · *θ_k_* i.e. the product of the disassembling rates of the two nodes.
2. Epidemic dynamics 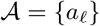:

- An infection emanating from an undiagnosed, infected individual 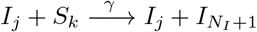 occurs with rate *γ* > 0 if node *j* and *k* are connected.
- An infection emanating from a diagnosed individual 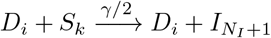 occurs with rate *γ* > 0 if node *j* and *k* are connected.
- An infected individual may be diagnosed with the infection 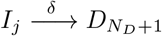, for all *j* = 1..*N_I_*.
- Individuals may die:

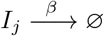, for all *j* = 1..*N_I_*
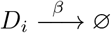, for all *i* = 1..*N_D_*

In many applications, we are only interested in the *epidemic dynamics*, e.g. to quantify the effect of particular epidemic control measures, such as contact restrictions or other behavioral changes, on the infection incidence. However, the transmission dynamics directly depend on the dynamics of the contact process. Moreover, an important characteristic of any *adaptive network* is that not only the contact dynamics affect the spreading process, but also vice versa (Fig. 1C). I.e. in the example above, the rate *λ_i_* and number of contacts of the individual depends on the actual state of node i. In our example, we sample 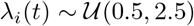 (1/day) and set *λ_i_*(*t*) = *λ_i_*(*t*) · 0.3 in case of a diagnosis.

Note that any time-dependent function is in principle possible (see *Discussion*).

The presented model can now be evolved using the stochastic simulation algorithm (SSA), Fig. 1D. The SSA would sample the time to the next event, which could either be an event that relates to the *contact dynamics*, or an event that relates to the *epidemic dynamics* as outlined in Algorithm 1.

#### Algorithm 1: SSA

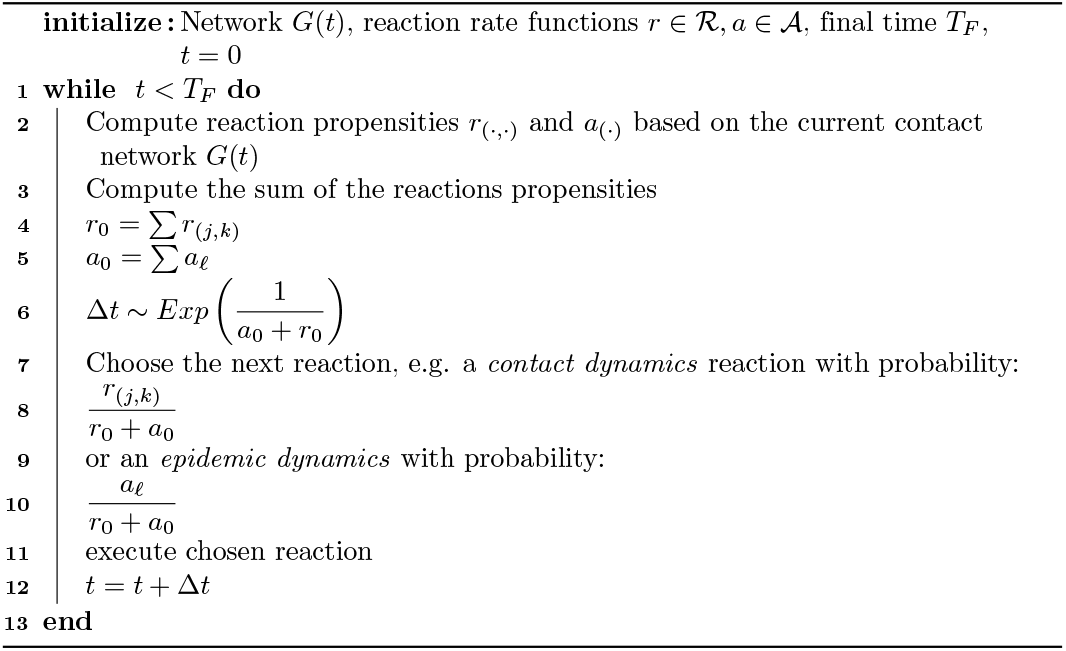

If the main interest is in assessing the *epidemic dynamics*, this approach would create considerable computational overhead, particularly if the *contact dynamics* are considerable faster than the *epidemic dynamics* (since only few contacts lead to infection, the *contact dynamics* are always faster than the *epidemic dynamics*).

In the following, we will present an algorithm that is able to precisely sample the time to the next infection event, while performing bulk updates on the *contact dynamics*. Notably, the bulk updates preserve all statistical properties of the *contact dynamics* that are relevant to the spreading process (*epidemic dynamics*). Hence, the algorithm allows to study the relevant dynamics at full resolution, while reducing computational burden.

### Simulation of effective spreading dynamics with SSATAN-X

In the proposed algorithm, we are splitting the above described stochastic process into a process *Y* comprising all reactions 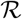 belonging to the *contact dynamics* and a process *Z* that is driven by reactions 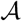 (*epidemic dynamics*). We then leap-forward the background process (Fig. 1E), until a reaction of the *epidemic process* happens (Alg. 2). Algorithm 2 implements the core idea of SSATAN-X and is an adaptation of the EXTRANDE method proposed by Voliotis in the context of Systems Biology [37].

#### Algorithm 2: SSATAN-X envelope algorithm for simulating effective spreading dynamics on adaptive networks.

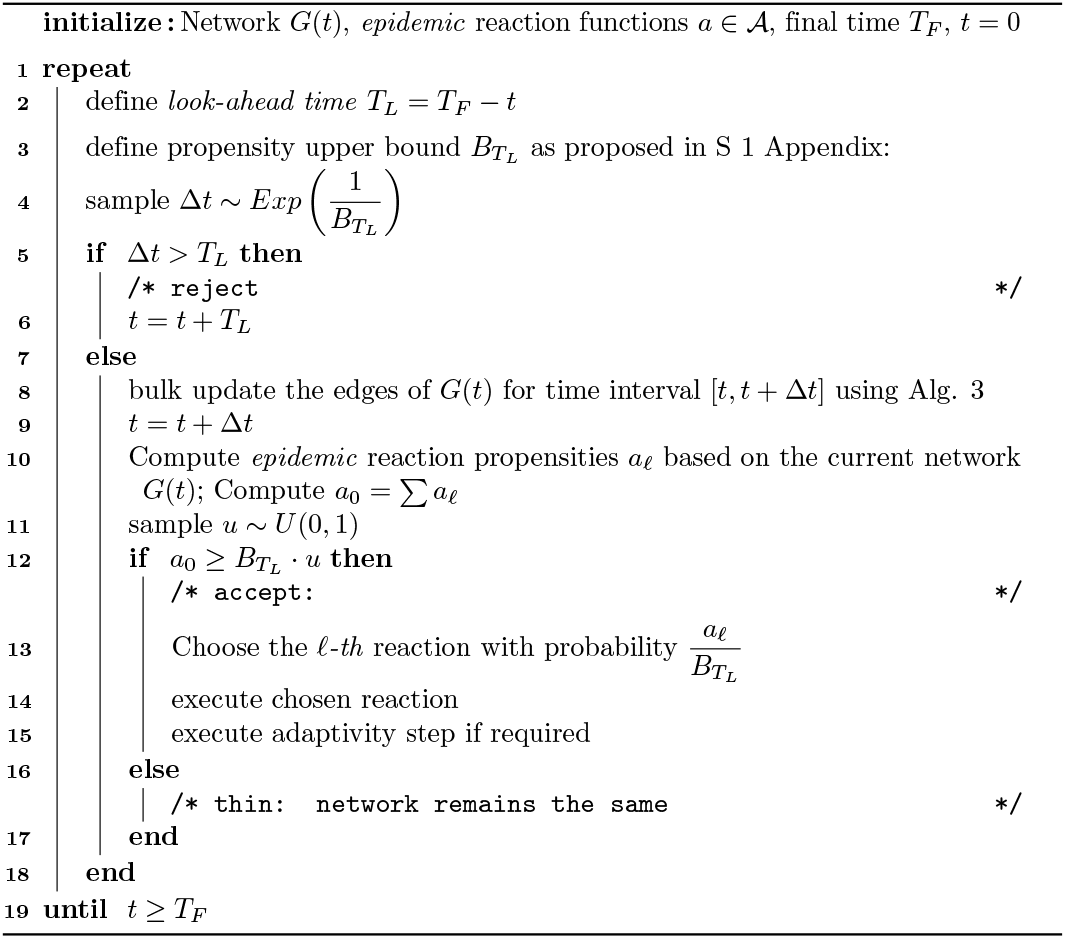

In the outline of the algorithm, **line 2** defines the so-called *Look-ahead* time *T_L_*, for which an upper bound of the sum of propensity functions *B_T_L__* ≥ *a*_0_ for the network *G*(*s*); *s* ∈ [*t; T_L_*] is calculated. According to [37] the *look-ahead* time horizon can safely be chosen as *T_L_* = *T_F_* – *t*.

Correspondingly, for *T_L_*, the upper bound *B_T_L__* of the sum of reaction propensities of the *main process* is calculated in **line 3**. The upper bound for the transmission reactions depends on the number of contacts (edges) between susceptible and diagnosed or undiagnosed individuals during *s* ∈ [*t; T_L_*]. As can be imagined, a large upper bound will result in many “thinning” reactions, which can make the algorithm inefficient. In S 1 Appendix, we provide a method to calculate *B_T_L__*. With respect to the upper bound *B_T_L__* we can propose the *time to the next reaction* 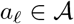, Δ*t* (**line 4**). Notably, since *B_T_L__* ≥ *a*_0_ for network *G*(*s*); *s* ∈ [*t; T_L_*], the proposed *time to the next reaction* might be shorter than the actual time to the next reaction event. This apparent under-estimation is “corrected” by the “thinning” step (**lines 16-17**), such that the statistics of the process are being preserved (for details refer to [37]). When the proposed time step Δ*t* lies within the look-ahead horizon *T_L_*, time *t* is updated by Δ*t* (**line 9**). At this point, the *actual* propensity functions *a_ℓ_* need to be updated. In order to do so, the contact dynamics (edges) of the network *G*(*t*) are bulk-updated using Algorithm 3 (tau-leaping)(**line 8**) and the next reaction to update (or “thin”) the state of the nodes (*main* process) is chosen (**lines 12–18**). Here, the adaptivity step (**line 15**) refers to any drastic changes to the contact network, when an *epidemic* reactions occurs. In our example, a diagnosed individual breaks all his/her contacts. The bulk-update (**line 8**) allows us to avoid the unnecessary computation of each individual contact update by reactions 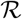 of the *contact* process.

### Bulk update of *contact network*

The bulk-updating algorithm is based on the *τ*-leaping algorithm, which was originally proposed by Cao and Gillespie [32] and was later modified by Anderson [38] by incorporating post-leap checks to guarantee accuracy.

We use the bulk-update algorithm only for reactions 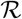 of the background process 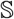 (contact dynamics). A key adaptation is to consider all reaction *of one type* ξ ∈ {+, −} (addition or deletion of edges) simultaneously. In the network example, only *two types* of reactions change the contact structure of the network: 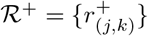 denotes the set of reactions that generate new edges {*v_j_, v_k_* }, *j* ≠ *k* with rate *λ_j_λ_k_*, whereas 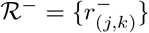 denotes the reactions that eliminate existing edges {*v_j_, v_k_*}, *j* ≠ *k* with rate *θ_j_θ_k_*. Hence, we have 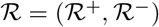.

In a finite population, the amount of edges in the network is limited by the number of edges in a full undirected graph *N*(*N* – 1)/2. Therefore, the number of edges available for deletion during one iteration is limited by the number of edges existing in the network |*E*(*t*)| at time *t*, such that the number of edges available for addition during one iteration is limited by *N*(*N* – 1)/2 – |*E*(*t*)|.

We denote a network *G* = {*E, V*}, as well as its complement network *G′* = {*E′, V′*}. To accurately capture the *contact dynamics*, we store both these networks. Both of the networks co-evolve synchronously (i.e. a deletion of an edge in the contact graph is accompanied by the addition of the same edge in the complement graph). This allows to apply less complex search algorithms to correctly simulate the contact dynamics. Adding and edge to *G* will be complemented by deleting this edge from *G′* and vice versa. Thus, we can describe the reactions of assembling and disassembling edges by the first order reactions:

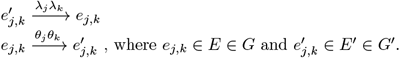

#### *τ*-leaping with post-leap checks

Algorithm 3 describes the *τ*-leaping algorithm, modified for bulk-updating of the *contact dynamics* of the network.

Based on the initial state of the system, the time step of the leap *τ* is calculated (**line 2**) based on the following formulas:

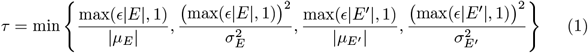

where |*E*|, |*E′*| denote the number of edges in the contact network and its complement graph. The *expected net* change to the network graph, as well as the complement graph is given by:

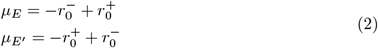

where 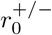 denote the sum of reaction propensities for *addition* and *deletion* respectively at the current network state *G*(*t*). The variance of the *net* change is given by

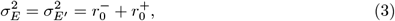

because both *addition* and *deletion* reactions are first-order reactions in the contact network and its complement graph.

##### Algorithm 3: *τ*-leaping for bulk-updating the contact network

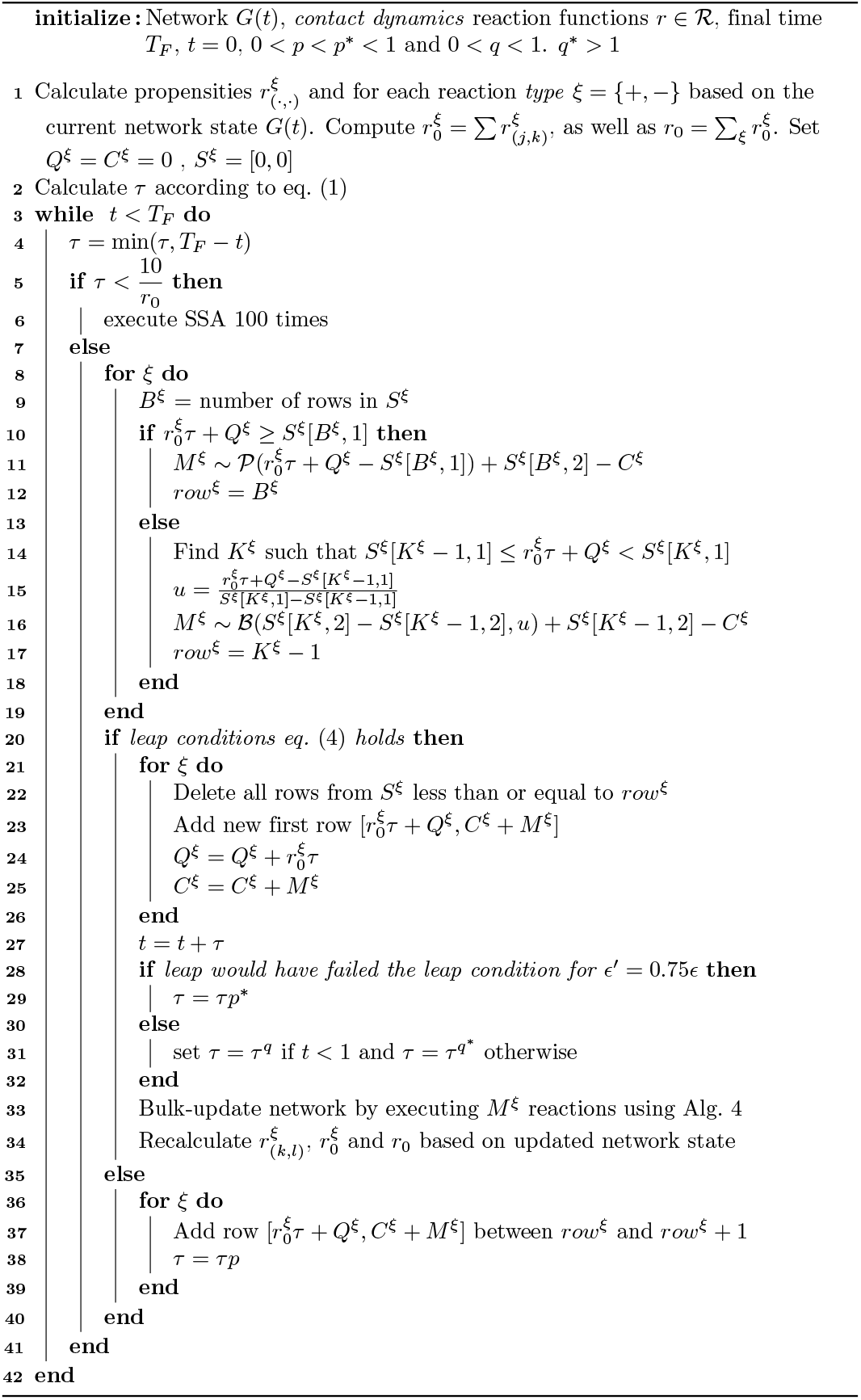

If the chosen *τ* is less than value of 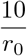, we abandon the leaping algorithm and execute a fixed amount of steps using the SSA algorithm (usually taken to be 100 [33 **lines 5–6**). Otherwise, for each reaction *type*, the number of executions *M^ξ^* is sample (**lines 9-19**), either from a Poisson distribution (**line 11**) or a Binomial distribution (**line 16**). If the leap is accepted (**line 20**) with condition

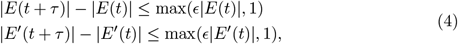

the time *t* is increased by *τ* (**line 27**), storage variables *T^ξ^, S^ξ^* and *C^ξ^* are updated (**lines 21-26**) and *τ* is updated (**lines 27-31**). The bulk update (**line 33**) is performed with respect with chosen values of *M^ξ^* and outlined in Alg. 4.

If the leap is rejected, the sampled values are stored (**lines 35-40**) and *τ* is decreased by multiplying it with 0 < *p* < 1. The algorithm returns to **line 4**.

##### Algorithm 4: Bulk Update

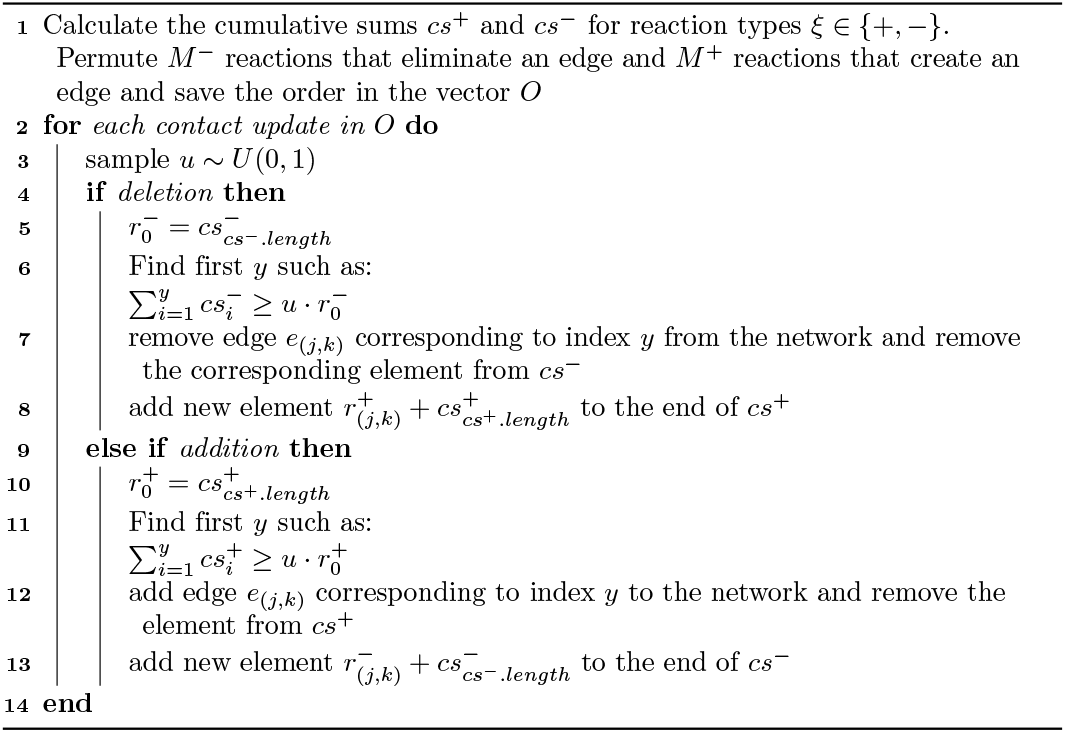

### Implementation and availability

We implemented the algorithm in C++ using an object-oriented approach. The contact network was implemented with the LEMON Graph library https://lemon.cs.elte.hu/trac/lemon

All codes are available from https://github.com/nmalysheva/SSATAN-X.

## Results

### SSATAN-X accurately captures contact network dynamics

In order to analyse the accuracy and performance of SSATAN-X, we first evaluated simulations with pure *contact dynamics*.

Whereas the exact stochastic simulation algorithm (Alg. 1) conducts changes to the contact dynamics network one-by-one, as illustrated in Fig. 2 (left), SSATAN-X uses bulk updates of the contact network, as described in Alg. 3 and shown in Fig. 2 (right). We remember that the main principle of SSATAN-X is that it accurately captures the statistics of the *contact dynamics* that is relevant to modelling the *epidemic dynamics*. Therefore, the statistics of the contact dynamics only need to be accurate at the time point when a bulk update was performed, but not in between bulk updates (Fig. 2, right).

**Fig 2.**
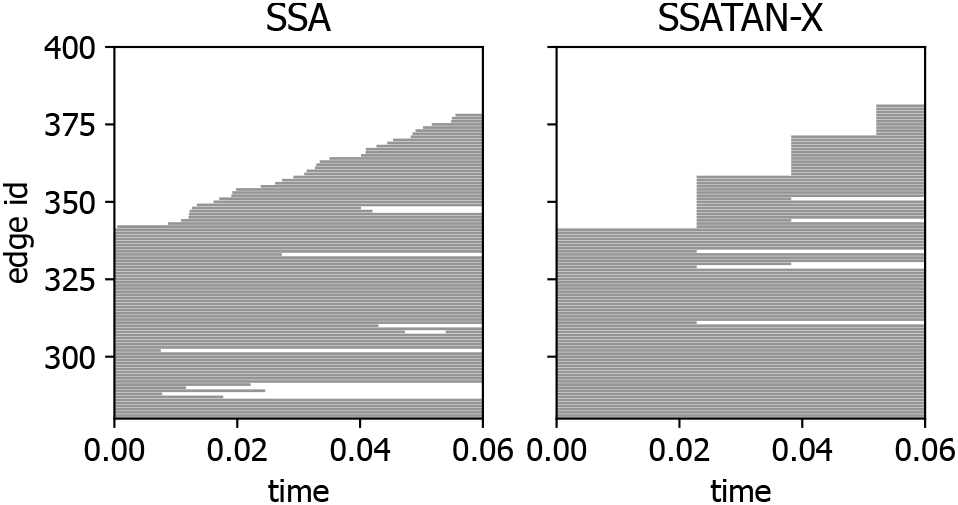
Edge activity plot. Edges of the network assigned to the id and depicted in the order of appearance. Each edge is drawn as a bar for the duration of its existence. **A.** SSA captures every single event (deletion or addition of an edge) individually, resulting in very small time steps. **B.** The bulk-updating in SSATAN-X allows to create and/or delete several edges using larger time steps, while maintaining the correct statistics at these discrete time steps.

To assess the accuracy of SSATAN-X in capturing the statistics of the *contact dynamics*, we first ran pure *contact dynamics* simulations, i.e. in which no epidemic transitions occur; *a_ℓ_* = 0 ∀ *ℓ*. We then compared 10^3^ trajectories of SSATAN-X with SSA simulations in time interval *t* ∈ [0, 5]. We initialized the population with 200 nodes and an initial amount of 3000 edges. Each node *j* was assigned a parameter 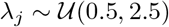 for establishing a new contact and 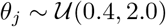 for loosing an existing contact.

Parameters for Algorithm 3 are set to be *p* = 0.75, *p*\* = 0.9, *q* = 0.98 [38], *ϵ* = 0.03 [32, 38], *q*\* = 1.02. Fig. 3 shows the degree distribution (mean ± standard deviation) of the contact network at different time points using SSA vs. SSATAN-X respectively. The simulation starts with a deterministic initial network, as can be seen by the absence of the shaded areas in Fig. 3A. During simulation, the degree distribution of the temporal network changes (Fig. 3B–D), until it reaches a stable distribution (Fig. 3E–F). During the entire simulation interval, both the mean degree distribution (solid blue vs. dashed red lines), as well as the standard deviations (blue and red shading) remain visually indistinguishable between the two algorithms.

**Fig 3.**
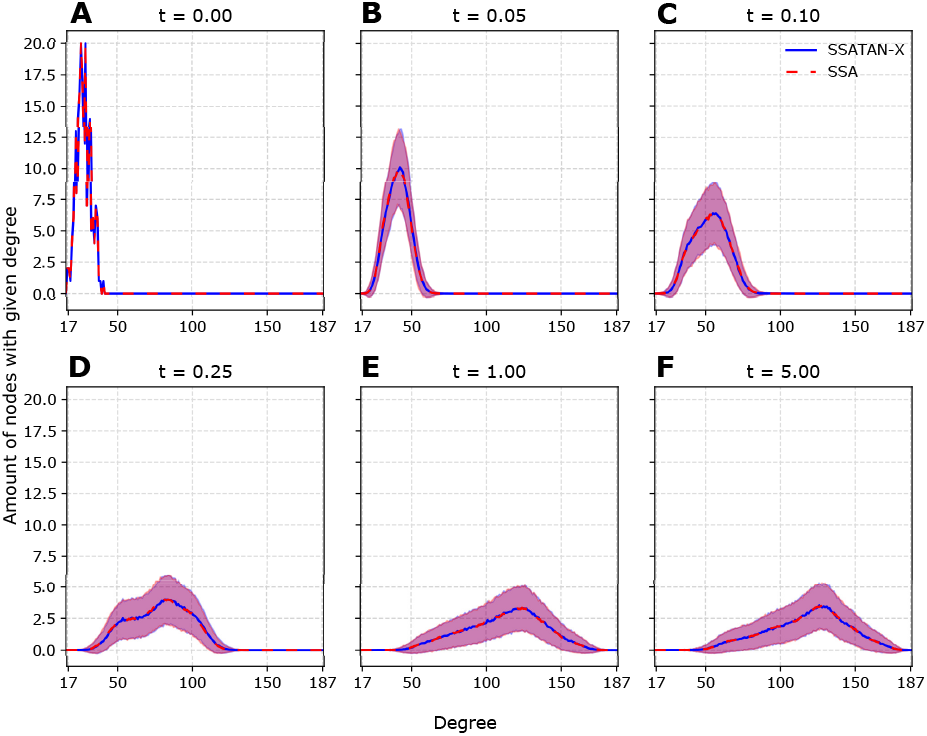
Degree distributions during simulation. The graphic depicts the mean degree distribution of the network during distinct time points within the simulation time (*t* ∈ [0, 5]). The dashed red and solid blue lines represent mean degree distribution over 10^3^ simulations using the SSA and SSATAN-X respectively. Shaded areas represent the corresponding standard deviations for the 10^3^ simulations.

To further quantify the differences between the evolving contact network over time, we computed the statistics of the Kolmogorov-Smirnov test, as shown in Fig. 4A–B. As can be seen in Fig. 4A, the probability that the degree distributions generated with the SSA vs. SSANTAN-X are identical is >> 0.95 (P-value) for all time points. The test statistic of the Kolmogorov-Smirnov test (distance between the respective ECDFs) is shown in Fig. 4B. The test statistic remained below the value of 0.0015 at all time points, further highlighting that differences in the degree distribution are insignificant.

**Fig 4.**
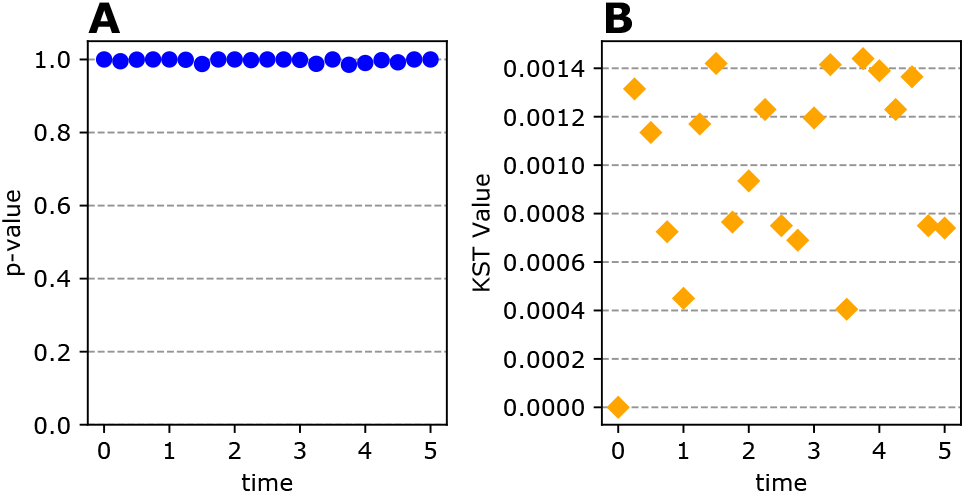
Statistical assessment of differences in the simulated contact network. We used the Kolmogorov-Smirnov test to compare the degree distributions obtained by SSA and SSATAN-X (represented on Fig. 3). **A.** The P-value denotes the probability that the two degree distributions derived from SSA- and SSATAN-X simulations respectively are identical. **B.** Statistics of the Kolmogorov-Smirnov test (KST), representing the distance between the two empirical cumulative distribution functions.

We hence conclude that the bulk-updating (Alg. 3) of the contact network dynamics, as implemented in SSATAN-X, accurately captures the statistics of the time-evolving contact network at the discrete time steps.

Next, we evaluate whether the SSATAN-X also accurately captures *epidemic dynamics* that unfold on an adaptive contact network.

### SSATAN-X accurately captures epidemic dynamics

In this section we present the results of the SSATAN-X algorithm applied to the adaptive susceptible-infected-diagnosed (SID) network model described in the *Methods* section with the following parameters: We initialize the *contact network* with 200 nodes (individuals) and 3000 edges connecting them. The *epidemic state* of the network is initialized with {S = 180, I=20, D=0}. For the *contact dynamics* each node *j* was assigned a parameter 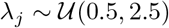 for establishing a new contact and 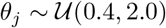 for loosing an existing contact. For the *epidemic dynamics*, the transmission rate along an S-I edge (contact) was set to *γ* = 0.04 and it was set to *γ*/2 along a S-D edge. The death rate of infected and diagnosed individuals was set to *β* = 0.08 and for susceptible it was set to *β* = 0. The diagnosis rate was set to *δ* = 0.5. In case of a diagnosis event, where the infected individual is becoming aware of his/her status, the newly diagnosed individual cuts all contacts and the individual’s rate of establishing new contact drops to 30% of the pre-diagnosis level.

In order to test the accuracy of the proposed algorithm, we conducted 10^3^ simulations using the SSA (Alg. 1), as well as SSATAN-X (Alg. 2-4). Foremost, we were interested to see whether the statistics of the times to the next *epidemic* event were correct, as proposed in Fig. 1D-E. In Fig. 5, we show a histogram of the time-steps to the next *epidemic* event (infection, diagnosis, death) for the SSA (orange bars), as well as SSATAN-X (blue bars). As can be seen, both algorithms yield indistinguishable statistics regarding the *epidemic* inter-event times. This reaffirms the correctness of the proposed algorithm in being able to modelling the *net* effect of contact dynamics on the spreading process. Fig. 6 shows the corresponding simulation results for the *epidemic* dynamics. The mean (± standard deviation) number of susceptible, infected and diagnosed are shown in Fig. 6A-C respectively. Both the mean and the standard deviation are indistinguishable when comparing simulations with the SSA (red line and red shading) vs. SSATAN-X (blue line and blue shading) over the entire duration of the simulation. Therefore, we can further confirm the correctness of SSATAN-X in simulating *epidemic dynamics* on an adaptive contact network.

**Fig 5.**
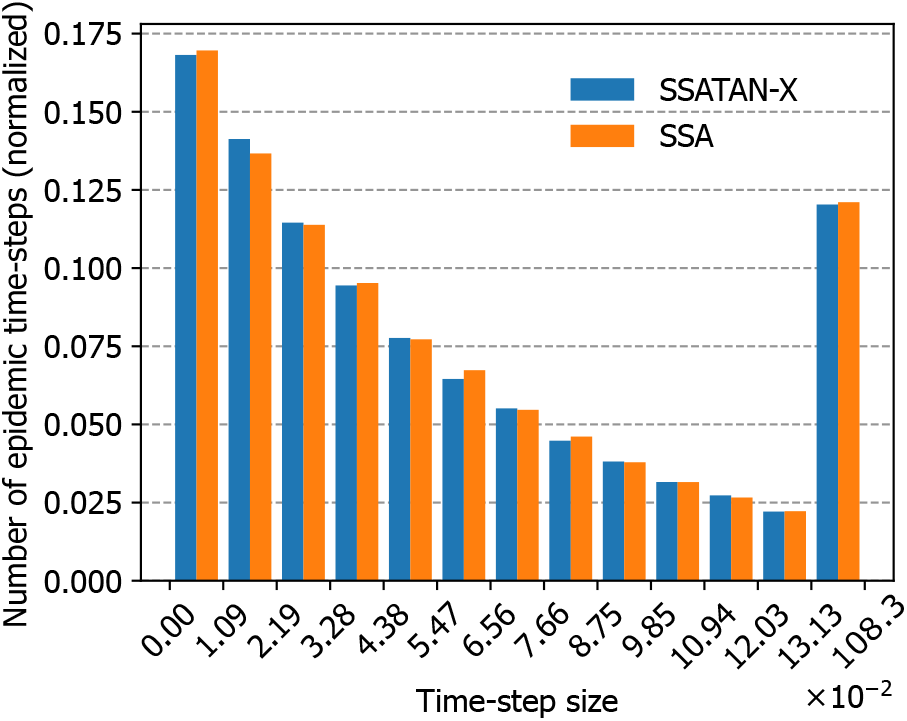
Time-step to the next epidemic event. The histogram depicts the frequency of the time-step sizes to the next epidemic event (infection, diagnosis or death), produced from 10^3^ simulations for each algorithm and normalized over the number of samples (overall 79375 *epidemic* time steps for SSATAN-X (blue bars) and 79025 for SSA (orange bars)).

**Fig 6.**
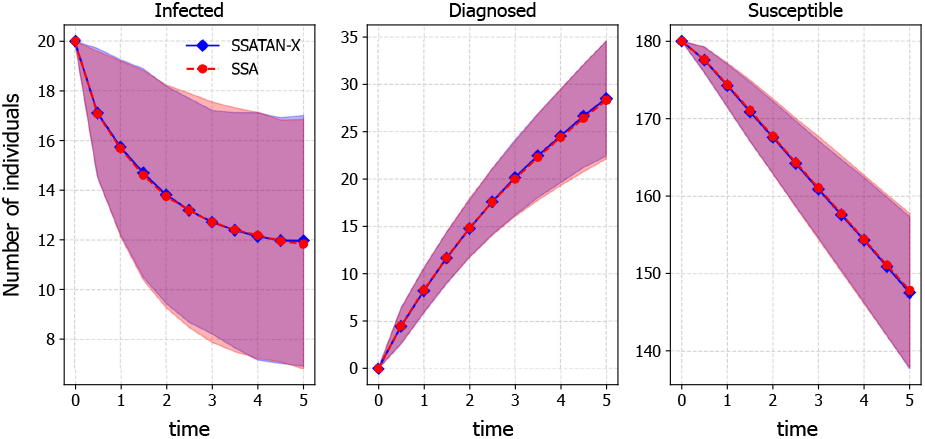
Simulated infection dynamics. The graphic depicts the change in the number of infected (panel **A.**), diagnosed (panel **B.**) and susceptible (panel **C.**) individuals over simulation time. Dashed red and solid blue lines describes the sample means over 10^3^ simulations using SSA and SSATAN-X respectively. Shaded areas represent the corresponding areas encompassed by the mean ± standard deviation.

### Impact of contact network adaptivity on the *epidemic dynamics*

The analysed S-I-D model (*Methods* section), belongs to the class of adaptive network models, because not only do the *contact dynamics* affect the *epidemic dynamics*, but *vice versa*, the *epidemic dynamics* also feed back onto the *contact dynamics* (compare Fig. 1C). Specifically, in the utilized model the *epidemic* event of diagnosis, changes both the contact network itself (cutting all edges of a newly diagnosed individual), but also its further dynamics (the rate of establishing new contacts drops to 30%).

To investigate the effect of adaptivity on the *epidemic*, as well as the *contact dynamics*, we consider two representative examples of the contact network model described above: for both examples, we start with {S = 180, I=20, D=0} and set the rates for establishing a new contact and for loosing an existing contact to 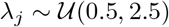 and 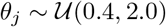, and the transmission rate to *γ* = 0.004. For the first of the examples we set the diagnosis rate to *δ* = 0, in other words, in this example we have a non-adaptive temporal network model (described in Fig. 1B). For the adaptive model (second example), we set the diagnosis rate to *δ* = 0.5. Adaptivity in the second example is implemented in the following way: The diagnosis event of an infected individual causes a complete rewiring of the individuals’ contact network. The individual looses all contacts upon diagnosis and subsequently has a decreased rate of making new contacts, i.e. *λ*_*j,t*+_ = 0.3 7 *λ*_*j,t*-_, where *t*^−^ and *t*^+^ denote the time before and after diagnosis. The transmission rate *γ* remains unchanged. Also, for both examples we set the death rate to *β* = 0. This allows to investigate the impact of adaptivity on the epidemic and contact dynamics, without the impact of other factors.

Fig. 7A shows the number of edges in the network that are eligible for transmission in the non-adaptive temporal network (*δ* = 0; blue error bars), vs. the adaptive network (*δ* = 0.5; orange error bars). Annotations show the percent of infected individuals that are diagnosed (e.g. D/(D+I)). Fig. 7B compares infection dynamics of both, the non-adaptive and adaptive model (blue vs. orange error bars). As can be seen, the diagnosis event alters the *epidemic dynamics* in the sense that less new infections occur in the adaptive network. In particular, since we set up the simulations in such way, that the transmission rates emanating from both infected and diagnosed were identical, the decrease in infection dynamics can be solely attributed to alterations in the *contact dynamics* in the adaptive networks (Fig. 7A), which then feed back onto the *epidemic dynamics*. I.e., the number of edges, eligible for the transmission of the virus decreases with an increasing percentage of infected individuals being diagnosed (Fig. 7A). The consequence are fewer opportunities for disease spreading and hence the number of new infections (orange error bar in Fig. 7B) decreases, too.

**Fig 7.**
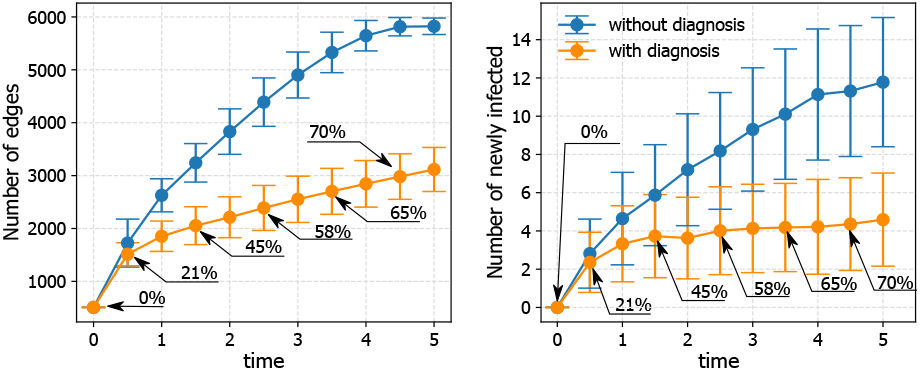
Impact of diagnosis on the degree distribution and amount of new infections. Blue error bars portray the results from a non-adaptive model, where we set the diagnosis rate *δ* = 0. Orange bars depict predictions from an adaptive model, where the diagnosis rate *δ* = 0.5. In the adaptive model, a diagnosis of individual *j*, leads to a loss of all contacts (self-isolation) and a reduced rate of forming new contacts *λ_j_* = 0.3 · *λ_j_*. Both panels depict results from 10^3^ SSATAN-X simulations for each parameter set. **A.** Error bars represent the mean amount ± standard deviation of the amount of edges eligible for transmission of the virus (i.e. number of contacts between infected and susceptibles and between diagnosed and susceptibles) over time. **B.** Error bars represent the mean amount ± standard deviation of newly infected individuals over time. Annotations on both panels show the percentage of infected individuals that are diagnosed (D)/(I + D).

### SSATAN-X speeds up computation time

Next, we analyzed the run time of SSATAN-X in comparison to SSA for different populations sizes (number of nodes). To this end, we took the model settings and parameters as described in section *SSATAN-X accurately captures epidemic dynamics*.

Simulations were started with fixed number of infected individuals {I=20}. The simulation results presented in Fig. 8A show that the run times increase with increasing population size. However, for the utilized parameter setting, the runtime of SSATAN-X is about 100-fold lower than the run time of SSA.

**Fig 8.**
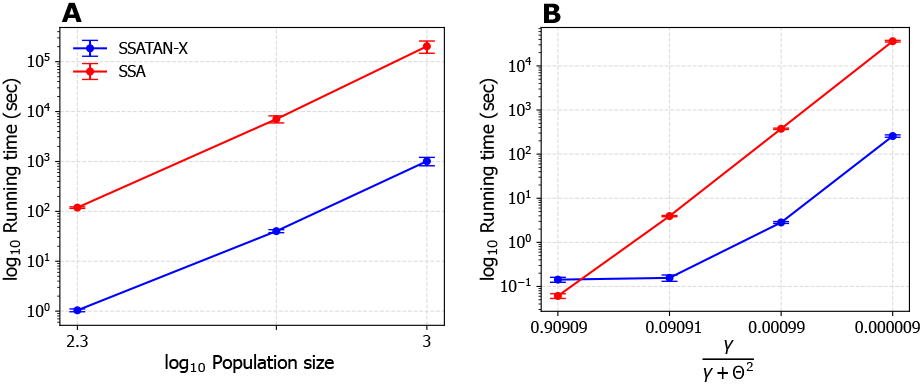
Run time performance. **A.** Log-log plot showing the mean (± standard deviation) run time of SSATAN-X (blue error bars) vs. SSA (red error bars) for different population sizes (200, 500 and 1000 individuals) over 10^3^ simulations for each parameter set on a single `2x AMD Epyc 7742 64-Core` compute node with a base clock of 2.25 Ghz respectively. **B.** Semi-log plot showing mean (± standard deviation) run time of SSATAN-X (blue error bars) vs. SSA (red error bars) for different parameterizations for a network with 200 individuals. The relation 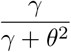, with *γ* being the transmission rate and *θ* being the rate of edge disassembling, can be interpreted as the probability of transmitting an infection before the contact is dissolving.

Next, we wanted to assess the runtime performance as a function of the utilized parameters. To that end, we chose homogeneous contact dynamics rates *λ_i_, θ_i_* across the population (all individuals having the same rates). This allows to assess how the run time changes as a function of the probability that transmission occurs on a contact edge before it dissolves. This probability is given by 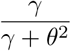, where *γ* is the transmission rate on a contact edge and *θ*^2^ is the rate of loosing an existing edge. In the analysis, we increased *λ* and *θ*.

The run time of both SSA and SSATAN-X for performing 10^3^ simulations for each parameter set is shown in Fig. 8B. As can be seen, when the relation 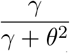 is close to 1, i.e. transmission is always bound to happen before an edge dissolves, SSA may be slightly faster than SSATAN-X. The reason is that the time-interval between two *epidemic* events becomes so small that the bulk update of the contact dynamics reduces to the SSA. Noteworthy, such models would actually not require to model separate *contact dynamics* altogether, making this example a bit artificial.

When the relation 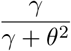 decreases, i.e. the *contact* dynamics being much faster than the *epidemic* dynamics, the run time of SSATAN-X becomes superior over SSA. Therefore, with low transmission rates and fast contact dynamics, SSATAN-X avoids any computational overheads from the *contact dynamics* and only takes their *net* effect on the *epidemiological dynamics* into account by *bulk updating*. On the contrary, SSA requires to execute each reaction (*contact* and *epidemic*) individually.

In summary, SSATAN-X is computationally more efficient than SSA. The run time of both algorithms scales with the population size and the “boost-factor” of SSATAN-X is preserved across different population sizes. However, the computational “boost-factor’ of SSATAN-X increases, when the *contact dynamics* are fast in comparison to the *epidemic dynamics*.

## Discussion

Spreading of infectious disease can occur when individuals encounter in contacts that are relevant for the route of transmission of a pathogen. These contacts can be sexual contacts for sexually transmitted disease, or spatial proximity for airborne infections. In order to more realistically model spreading of infectious diseases, different network models have been proposed in the past [10]. *Adaptive networks* encompass a class of models, where a contact network structure changes dynamically in response to the dynamics of the spreading process [9]. These types of models may therefore allow to realistically consider behavioural changes that occur when individuals become aware of their infection and self-isolate [39], or experience debilitating symptoms.

In this article, we introduce SSATAN-X, an efficient and exact algorithm to simulate effective spreading dynamics on adaptive networks.

SSATAN-X takes advantage of the fact that *contact dynamics* must be faster than *epidemic dynamics*. Instead of performing detailed simulations of the underlying contact dynamics, the core idea of SSATAN-X is to only consider the effect that the underlying *contact dynamics* have on the *epidemic dynamics*. For this purpose, SSATAN-X uses bulk updates of the contact structure between *epidemic* events, such as disease transmission, diagnosis or death (Fig. 2).

In Figure 3, we show that the proposed bulk-updating conserves the statistical properties of the underlying contact network at times that are relevant to the *epidemic dynamics*. Because the statistics of the underlying contact network at times that are relevant to the *epidemic dynamics* are preserved, the statistics related to the firing of the next epidemic event (Fig. 5) are preserved, too. Consequently, the resulting *epidemic dynamics* are statistically correct, Fig. 6–7. However, the bulk updating avoids computational overheads from updating contact edges one-by-one. Therefore, depending on the choice of parameters, SSATAN-X can speed up computation time quite significantly in comparison to exact methods like the stochastic simulation algorithm (SSA) without any loss in simulation accuracy, Fig. 8. The speed up factor critically depends on the relation of the *contacts dynamics* to the *transmission dynamics*. For example, if contacts generally do not result in transmission, computational speed up with SSATAN-X will be considerable. On the other hand, if transmission will always occur on a contact edge before it dissolves, the computational speed up will only be moderate or even absent. In Fig. 8B, we expressed this dependency as *γ*/(*γ* + *θ*^2^), where *θ*^2^ denotes the rate of loosing an edge and *γ* is the transmission rate along a contact edge.

Note that the transmission rate *γ* denotes a lumped parameter, that can be further decomposed to allow for more sophisticated modelling, realistic parameterization and model extensions: For example, if exposures on a contact edge occur with rate r_e_ and if the exposures are of the same type, then the transmission rate is just a scaled variant of the exposure rate, i.e. *γ* = *r_e_* · *p*_tr|*e*_, where *p*_tr|*e*_ is the probability that a transmission occurs for a single exposure event e. Notably, for many infectious diseases and types of exposures *p*_tr|*e*_ is readily known [25–27] and the effects of pharmaceutic [30, 31] or non-pharmaceutic interventions [40, 41] on *p*_tr|*e*_ can be quantified, allowing for model extensions that can link within-host pathogen dynamics to the spreading dynamics at the population level.

Taken together, the lumped transmission rate *γ* is a function of the transmission probability per exposure event *p*_tr|*e*_, as well as the frequency of exposures r_e_ when individuals are in contact. For many infectious diseases and types of contacts it may be argued that *γ*/(*γ* + *θ*^2^) << 1; I.e., most transient contacts may not result in pathogen transmission. Behavioral changes, such as distancing, isolation or quarantine, may further reduce the transmission rate by reducing r_e_, whereas pharmaceutic (vaccination, prophylaxis, treatment) [42–44] or non-pharmaceutic interventions (e.g. mask wearing for airborne-, and condom usage for sexually transmitted infections) [45, 46] may lower *p*_tr|*e*_ in a static or time-dependent manner.

In our examples, we considered reaction rates (for 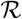 and 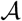) to be constants. However, it is also possible to consider time-dependent rates: For example, the transmission rate *γ* may be a function of time, due to some time-dependent pharmacokinetic effect of a prophylaxis [28, 29, 47], or depending on how long an individual is being infected or contagious [40]. These time-dependent effects may be incorporated when determining the upper bound B in Algorithm 2 (S 1 Appendix). While this modelling may further reduce the value of B it may even boost the overall computational performance by increasing the time to the next epidemiological event.

In our paper, we use the Anderson tau-leap algorithm (Alg. 3) [36] to compute the bulk updates of the *contact dynamics* between two epidemiological events. It is also possible to use other tau-leap based approaches, as summarized in [48], however, we used the Anderson tau-leap algorithm, because it does guarantee accuracy through step size adaptation.

While *τ*–leaping may be considered as a stochastic simulation algorithm that has some analogies to explicit (first order) Euler methods for solving systems of ordinary differential equations (ODE) [49], another possibility for the bulk updating of the *contact dynamics* is to adapt higher-order schemes with adaptive step sizes that have been proposed in the context of ODEs [50]. For example, the Runge-Kutta-Fehlberg method [51] could be used to adapt step sizes for the bulk updating. A clear advantage of higher-order schemes in the context of bulk-updating the contact network is the fact that larger step sized may be chosen that preserve accuracy [52]. In the future, we will further improve SSATAN-X by incorporating these higher-order approximation schemes.

In our paper we consider finite populations. In finite populations, the maximum possible amount of contacts of the each individual *i* has an upper theoretical limit of *N* – 1 and a complete graph would have *N* · (*N* – 1)/2 edges. This gives us a straightforward way to estimate the total number of edges available for deletion or addition at any time point t: The number of edges available for deletion equals the number of edges present *E*(*t*), whereas the number of edges available for addition is given by *N* · (*N* – 1)/2 – *E*(*t*) or in other words, the number of edges in a complement graph. Note that for infinite populations the number of edges that can be deleted is still given by *E*(*t*), whereas, for adding edges one may simply abandon the ‘complement graph’ approach, making the algorithm more efficient.

Some authors have also proposed more restrictive models for the number of contacts [11]. Some of these concepts may be inspired by the popular Dunbar number [53, 54], which is rooted in the presumption that individuals may not maintain stable social relationships with more than 150 other individuals. Notably, Dunbar’s presumption has not been without critique [55].

However, contact networks may exist, in which the maximal number of contacts of an individual is less than the number of available individuals in the network. For example, Schmid and Kretzschmar [11] describe an individual-based model, in which each individual has an upper-limit capacity regarding the number of concurrent sexual partnerships, with pair formation and separation simulated as a dynamic processes, where each individual has his/her own probabilities of contact formation and separation drawn from an appropriate distribution. To model these kinds of restrictive contact dynamics with SSATAN-X, the number of edges available for deletion remains unaffected, while the estimation of the number of edges available for addition becomes more involved: Depending on the order of the addition and remaining contact capacities, there may be several outcomes with the different amounts of edges possible to be added. For example, in a network where all nodes but one have exhausted their capacities, no more edges can be added without violating the contact constraints. Therefore, it becomes necessary to estimate a “worst scenario” case, i.e. a contact bulk update that resulted in the least overall number of edges. To do this, the Havel-Hakimi algorithm [56] with modifications that consider existing edges *E*(*t*) may be employed.

## Conclusion

We introduced SSATAN-X, an efficient stochastic algorithm to simulate spreading processes on adaptive networks. The core idea of SSATAN-X is to perform bulk updates of *contact dynamics*, to accurately estimate their net effect on the *epidemic* dynamics. SSATAN-X may be extended to allow for more detailed modelling of interventions on pathogen transmission, as well as regarding the contact dynamics. The *epidemic dynamics* are propagated using EXTRANDE, while the bulk updating of *contact dynamics* has been implemented using Anderson-Tau leaping [38]. However, any method that preserved the statistical properties of the underlying contact network may, in principle, be used.

## Supporting information

**S1 Appendix. Determination of upper bound B.** In the Supplementary text, the determination of the upper bound for the sum of propensities of the *epidemic dynamics* is described.

## Acknowledgments

We thank Benjamin Maier and Dirk Brockmann for fruitful discussions. MvK and NM acknowledge funding from the German Ministry for Science and Education (BMBF; grant numbers 031A307 and 01KI2016). NM acknowledges funding provided through the International Max-Planck Research School “Biology and Computation” (IMPRS-BAC). The funders had no role in designing the research or the decision to publish.

## Notes

### Competing Interest Statement

The authors have declared no competing interest.

